# Adaptive laboratory evolution triggers pathogen-dependent broad-spectrum antimicrobial potency in *Streptomyces*

**DOI:** 10.1101/2021.01.20.427378

**Authors:** Dharmesh Harwani, Jyotsna Begani, Sweta Barupal, Jyoti Lakhani

## Abstract

In the present study, adaptive laboratory evolution was used to stimulate antibiotic production in a weak antibiotic-producing *Streptomyces* strain JB140. The seven different competition experiments utilized three serial passages (three cycles of adaptation-selection of 15 days each) of a weak antibiotic-producing *Streptomyces* strain (wild-type) against one (biculture) or two (triculture) or three (quadriculture) target pathogens. This resulted in the evolution of a weak antibiotic-producing strain into the seven unique mutant phenotypes that acquired the ability to constitutively exhibit increased antimicrobial activity against bacterial pathogens. The mutant not only effectively inhibited the growth of the tested pathogens but also observed to produce antimicrobial against multidrug-resistant (MDR) *E. coli.* Intriguingly, the highest antimicrobial activity was registered with the *Streptomyces* mutants that were adaptively evolved against the three pathogens (quadriculture competition). In contrast to the adaptively evolved mutants, a weak antimicrobial activity was detected in the un-evolved, wild-type *Streptomyces*. To get molecular evidence of evolution, RAPD profiles of the wild-type *Streptomyces* and its evolved mutants were compared that revealed significant polymorphism among them. These results demonstrated that competition-based adaptive laboratory evolution method can constitute a platform for evolutionary engineering to select improved phenotypes (mutants) with increased production of antibiotics against targeted pathogens.

## Introduction

The discovery of antibiotics by Alexander Flemming in 1929 revolutionized the treatment of diseases caused by microbial pathogens (Demain and Elander 1999). At present times, the numbers of infection caused by MDR *Staphylococcus aureus, Acinetobacter baumannii, Klebsiella pneumoniae, Pseudomonas aeruginosa* and *Mycobacterium tuberculosis* have been found to pose serious threats to human health. Consequently, efforts are being made globally to develop new antibiotic compounds to antagonize these deadly pathogens (Appelbaum 2006; Boucher et al. 2009). The majority of antibiotics are natural products or their semi-synthetic derivatives that originated from the genus *Streptomyces* in the order *Actinomycetales* (Watve et al. 2001; Newman et al. 2007), that are known to be distributed in soil and marine sediments. The clinically important antibiotics produced by *Streptomyces* species include tetracyclines, aminoglycosides (streptomycin, neomycin, kanamycin), macrolides (erythromycin), chloramphenicol and rifamycins (Berdy 2005; Demain and Sanchez 2009). Many new antibiotics were isolated from actinomycetes between the late 1940s and the late 1960s, a period is known as the “Golden Age” of antibiotic discovery. After that, the rate of discovery gradually declined due to the frequent re-discovery of already known chemical compounds. However, the recommendations from the recent genome sequence information confirm that this source is still not yet exhausted (Bentley et al. 2002; Ikeda et al. 2003; Medema et al. 2010; Ohnishi et al. 2008; Song et al. 2010; Jiao et al. 2018).

Many attempts have been made by researchers to develop methods to stimulate the production of new chemical molecules (Mutka et al. 2006; Gottelt et al. 2010; Scherlach et al.2010; Laureti 2011; Baltz et al. 2011; Garbeva et al. 2011). One of these methods includes adaptive laboratory evolution that has been successfully used to augment the production of antimicrobial molecules (Goers et al. 2014; Charusanti et al. 2012; Stergiopoulos et al. 2013; Blum et al., 2016; Chunxiao et al. 2018). The adaptive laboratory evolution is a phenomenon in which a parent strain is serially passed against a certain selection pressure for few generations to promote adaptation to a new prevailing niche. In addition, this phenomenon has also helped to increase our current understanding of natural laws of evolution that may further provide convincing solutions to the rising problems of multidrug resistance (Cohen et al. 1989; Donald and van Helden 2009; Wensing et al. 2015) and cancer (Riganti et al. 2015). In the present paper, we speculated that a weak antibiotic strain of *Streptomyces* could undergo evolution after getting adapted in the laboratory if challenged against a bacterial pathogen. Because of serial passages against the challenging pathogen, it may now produce antimicrobials that are not normally produced by its wild-type parental strain. It was also hypothesized that challenging the wild-type strain with more than one pathogen in adaptive evolution may elicit a different response.

In other studies, co-cultures of two different organisms have been observed to improve the production of antimicrobial molecules (Slattery et al. 2001; Oh et al. 2007; Kurosawa et al. 2008; Bills et al. 2009; Garbeva et al. 2011; Harwani et al. 2018). Importantly, these experiments have not used adaptive-selection cycles as these are used in the present study. The co-culture experiments involve the cultivation of two (or more) microbes (interspecies interactions) in the same closed and restricted environment for a certain period. The supernatant from these mixed cultures and mono-culture (control) are then analyzed for their antagonistic potential. If no bio-activity is detected, the experiment is considered to be over and not carried forward. While as compared to the co-culture, in the laboratory evolution experiments described in the present study, the wild-type strain (here *Streptomyces* strain JB140) is co-cultured with different bacterial pathogens in the bi, tri and quadriculture experiments. After purification of the *Streptomyces* phenotypes from the first cycle, from the mixed cultures, the experiments involving serial passages of the selected phenotypes against bacterial pathogens are continued until the antagonistic potential is detected. In this way, the present method utilized three serial passages (three successive cycles of adaptation-selection of 15 days each) of JB140^WT^ against bacterial pathogens. Consequently, it led to the identification of seven unique mutants that were found to produce an increased amount of antimicrobials to inhibit the growth of bacterial pathogens effectively, including MDR *E. coli*, as compared to their parental wild-type counterpart. Interestingly, when the challenge was posed by more than one pathogen in the laboratory evolution experiments, the stronger antagonistic response was elicited by the producer strain. Thus the requirement of three adaptation-selection cycles was crucial for the wild-type producer strain to exhibit stimulated production of antimicrobials against the target pathogens. Besides this, we could be able to collect the molecular evidence of the adaptive laboratory evolution when RAPD profiles of the wild-type *Streptomyces* and its evolved mutants were compared.

## Material and Methods

The present research utilized a weak antibiotic-producing, *Streptomyces* strain JB140 (wild-type), its seven mutants namely JB140^*Pv*^, JB140^*St*^, JB140^*Sa*^, JB140^*Sa,St*^, JB140^*Sa,Pv*^, JB140^*St,Pv*^, JB140^*Sa,St,Pv*^, three bacterial pathogens *Salmonella typhi* (NCIM 2051), *Staphylococcus aureus* (NCIM 2079), *Proteus vulgaris* (NCIM 2027) and one MDR clinical isolate *Escherichia coli.*

### Selective isolation, maintenance and characterization

The streptomycete isolate JB140 was isolated from the soil sample collected at Ganganagar (29.9038°N, 73.8772°E) region falling in the arid Thar desert in the Indian state Rajasthan. After selective isolation, using the <u>*M*</u>odified <u>*A*</u>ctinomycete <u>*S*</u>elective (MAS) agar medium, the pure culture of the isolate was stored at −40°C using 20% (w/v) sterile glycerol until further use. The isolate JB140 was further subjected to its morphological studies using the bright field and scanning electron microscopy (SEM). The sample for SEM analysis was prepared according to the protocol described by Srinivasan et al. (2014). Phenotypic characterization of the isolate was performed by growing it on ISP (International *Streptomyces* Project) media as described by Shirling and Gottlieb (1966).

### Fermentation and evaluation of antimicrobial activity of the isolate JB140

After phenotypic characterization, the isolate was screened for its antibacterial activity against bacterial pathogens in primary and secondary screening using agar plug and agar well diffusion methods respectively. For small scale fermentation, the isolate was cultured in a 1L flask containing 500mL of ISP2 medium. The fermentation was carried out at 40°C for 10 days at an agitation rate of 120rpm. The growth was monitored by measuring optical density at 600nm for 15 days at two days interval. After cultivation, the fermentation broth was extracted with ethyl acetate. The extract was concentrated to dryness and the residue crude extract was dissolved in methanol and stored at 4°C for further use. The methanolic extracts were used to evaluate antimicrobial activity in the agar well diffusion method. During the time course of fermentation, antimicrobial activity at different time points was also monitored. The antimicrobial activity was assayed against three type strains *Salmonella typhi* (NCIM 2051), *Staphylococcus aureus* (NCIM 2079), *Proteus vulgaris* (NCIM 2027) and one MDR clinical isolate *Escherichia coli.*

### Adaptive evolution protocol

In the present adaptive laboratory evolution protocol, seven different competition experiments were performed in which a weak antibiotic-producing, wild-type (WT) *Streptomyces* strain JB140 was challenged with bacterial pathogens in there different ways as depicted in figure 1. The first three (1^st^, 2^nd^, 3^rd^) experiments involved JB140^WT^ and one of the three target pathogens (a bi-culture competition consisted of JB140^WT^ and *P. vulgaris* or *S. typhi* or *S. aureus).* The next three experiments (4^th^, 5^th^, 6^th^) involved JB140^WT^ and two target pathogens (a tri-culture competition consisted of JB140^WT^ and *S. aureus+S. typhi* or *S. aureus+P. vulgaris* or *S. typhi+P. vulgaris).* The last experiment (7^th^) involved JB140^WT^ and three target pathogens (a quadri-culture competition consisted of JB140^WT^ and *S. aureus+S. typhi+P. vulgaris).* A mono-culture involving only JB140^WT^ served as an internal control. The seven individual experiments were performed by inoculating 50mL of ISP2 broth medium in a 250mL flask with a loopful of JB140^WT^ mycelia. After inoculation, the flasks were covered with cotton plug and incubated at 37°C in a shaker incubator at 100rpm for 48h. After observing the visible growth of JB140^WT^, flasks for bi-culture, tri-culture and quadri-culture were inoculated with exactly 100 microlitres each (optical density at 600nm=0.50) of the target pathogens accordingly and incubated at 37°C for ten days as shown in figure 1 (step 1).

**Fig. 1.**
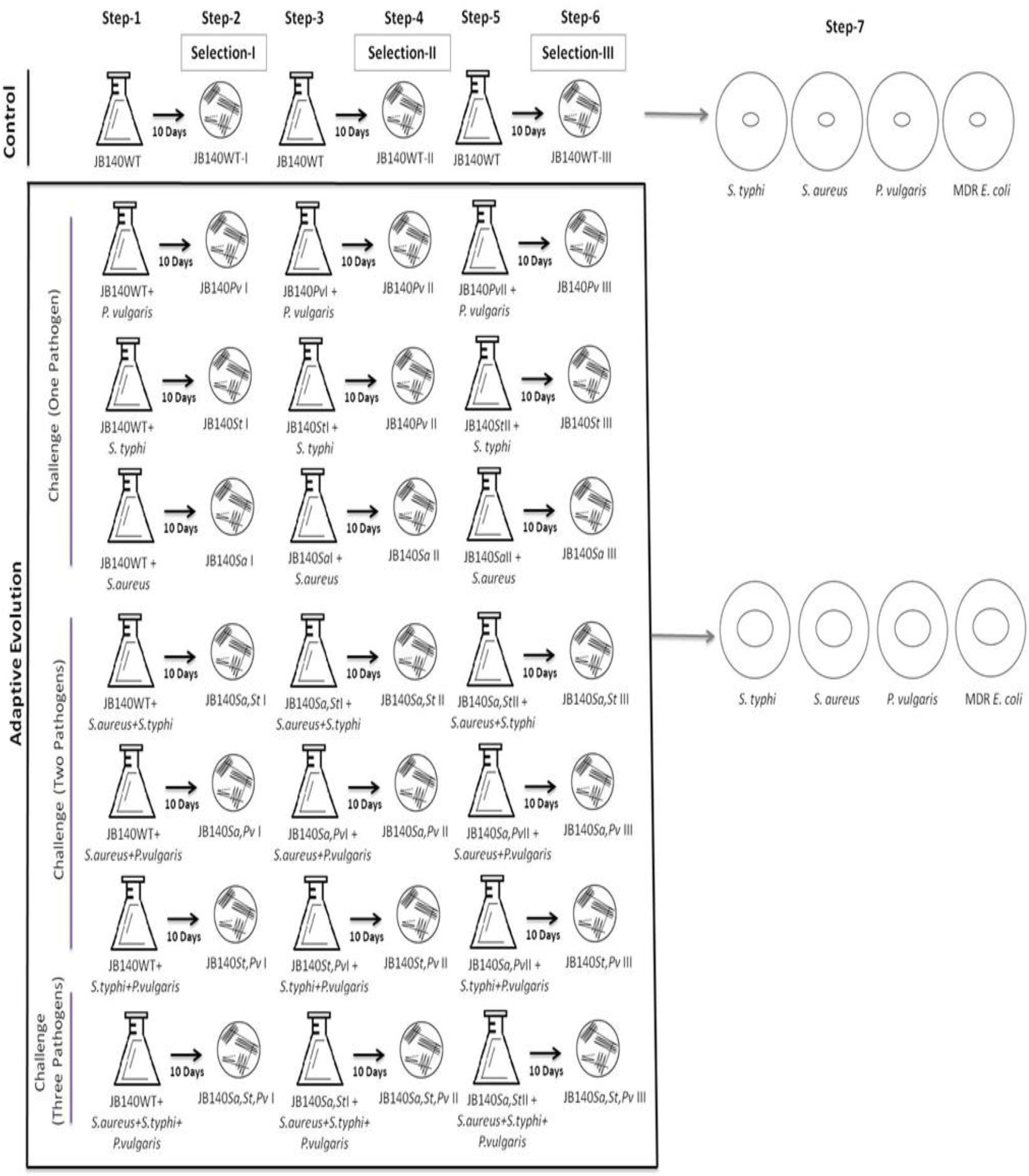
Adaptive evolution protocol (seven-steps) wherein a weak antibiotic-producing *Streptomyces* strain JB140 was competed against bacterial pathogens to stimulate antibiotic production

The flask for monoculture contained only JB140^WT^. After this step, on the eleventh day, 100 microlitres from ten days old mono-culture, bi-culture, tri-culture and quadri-culture were spread on the fresh ISP2 agar medium and incubated at 37°C for five days to separate JB140^WT-^I (control), JB140^*Pv-*^I (bi-culture), JB140^*St-*^I (bi-culture), JB140^*Sa-*^I (bi-culture), JB140^*Sa,St-*^I (triculture), JB140^*Sa,Pv-*^I (tri-culture), JB140^*St,Pv-*^I (tri-culture), JB140^*Sa,St,Pv-*^I (quadri-culture) from their respective flasks (step 2) (Fig. 1). After purification, the next cycle of adaptation-selection was repeated by inoculating purified wild-type and evolved mutants from step 2 into the fresh ISP2 medium. It is important to mention here that the flasks in the successive steps were introduced with freshly grown pathogen/s from their respective stocks. The inoculated flasks were incubated for another ten days (step 3; II cycle). After incubation, the mixed cultures were purified to obtain JB140^WT-^II (control), JB140^*Pv-*^II (bi-culture), JB140^*St-*^II (bi-culture), JB140^*Sa-*^II (bi-culture), JB140^*Sa,St-*^II (tri-culture), JB140^*Sa,Pv-*^II (tri-culture), JB140^*St,Pv-*^II (tri-culture), JB140^*Sa,St,Pv-*^II (quadri-culture) from their respective flasks (step 4). The adaptive evolution cycle (III cycle) was again repeated to start a new adaptation and selection cycle in step 5 and 6 to finally receive the wild-type JB140^WT-^III and evolved mutants JB140^*Pv-*^III, JB140^*St-*^III, JB140^*Sa-*^III, JB140^*Sa,St-*^III, JB140^*Sa,Pv-*^III, JB140^*St,Pv-*^III, JB140^*Sa,St,Pv-*^III.

The protocol described here, consisted of six main steps that took ~45 days to complete (Fig. 1). The wild-type and evolved isolates were preserved in the glycerol stocks (20%v/v) at −40°C. After each cycle of adaption-selection, the antimicrobial activities of the wild-type and evolved mutants were assayed against pathogenic *Salmonella typhi* (NCIM 2051), *Staphylococcus aureus* (NCIM 2079) and *Proteus vulgaris* (NCIM 2027) using agar well diffusion method. The antimicrobial activity of the mutants purified after the third cycle was also tested against MDR *Escherichia coli* (clinical isolate responsible for <u>*U*</u>rinary <u>*T*</u>ract <u>*i*</u>nfection (UTI)) in step 7. The detailed prototype of the present method has been also provided in supplementary data (Fig.S1 and Fig. S2).

### DNA extraction, sequencing and phylogenetic analysis

The wild-type isolate JB140 and evolved mutants JB140^*Pv-*^III, JB140^*St-*^III, JB140^*Sa-*^III, JB140^*Sa,Sp*^ III, JB140^*Sa,Pv-*^III, JB140^*St,Pv-*^III, JB140^*Sa,St,Pv-*^III were subjected to 16SrRNA gene sequence analysis for their precise identification. The genomic DNA of each isolate was extracted using a method of Li et al. (2007). 16SrRNA gene from each was amplified using a primer pair 27F 5’AGA GTT TGA TCC TGG CTC AG3’ and 1492R 5’GGT TAC CTT GTT ACG ACT T3’. The reaction started with an initial denaturation at 95°C for 5min, followed by 30 cycles of DNA denaturation at 95°C for 1 min, primer-annealing at 44°C for 1 min and extension cycle at 72°C for 1.5 min, with a final extension at 72°C for 10 min. PCR-amplicons were visualized in 2% agarose gel electrophoresis and subsequently revealed with ethidium bromide staining. The amplified 16SrDNA gene product was sequenced using Sanger didoxy method. Manual sequence edition, alignment, and contig assembly were performed using Vector NTI v10 software package. Sequencing results were analyzed for chimeras using DECIPHER v1.4.0 program (Wright et al. 2012). The 16SrRNA gene sequences were compared with the GenBank/EMBL/DDBI databases by using the BLASTN (Altschul et al. 1997) search program. After pairwise alignment using CLUSTAL_X program v1.8 (Thompson et al. 1997), the phylogenetic tree of the wild-type isolate JB140 was constructed by neighbor-joining (Saitou and Nei 1987) method using MEGA X (Kumar et al. 2018). The evolutionary distance was computed using the Kimura 2-parameter method (Kimura 1980). The stability of relationships was assessed by performing bootstrap analyses (Felsenstein 1985) of the neighbor-joining data based on 1,000 re-samplings.

### RAPD profiling

Random amplified polymorphic DNA (RAPD) analysis of the wild-type strain JB140 and evolved mutants was performed by using OPERON random primers (USA); RAPD-1 5’AAGAGCCCGT3’, RAPD-2 5’GTTTCCGCCC 3’ and RAPD-3 5’GAGGCCCTTC 3’. These three RAPD primers were used individually or in combination (all three) in separate amplification cycles. The reaction started with an initial denaturation at 95°C for 5min, followed by 40 cycles of DNA denaturation, at 95°C for 1 min, primer-annealing at 32°C for 1 min and extension cycle at 72°C for 1 min, with a final extension at 72°C for 10 min. PCR-amplicons were visualized in 2% agarose gel electrophoresis after ethidium bromide staining. The detailed prototype of the present method has been also provided in supplementary data (Fig. S2).

## Results

### Isolation and characterization of JB140

Based on characteristic colonial morphology, notably the ability to form aerial hypha and substrate mycelia, earthy smell and chalky appearance of the mature colony, darker in the center and lighter farther, irregular, fuzzy edge, pigmented, strongly adhered and leathery texture, the isolate was putatively identified as actinobacteria. The Gram-positive actinobacterial isolate JB140 showed typical mycelial structure in Gram’s staining (Fig. 2A). The scanning electron micrograph of the isolate JB140 revealed that it has a long looped retinaculiaperti spore chain with a smooth surface (Fig. 2B). The isolate exhibited well developed aerial and substrate mycelium on various ISP media (Fig. 3).

**Fig. 2.**
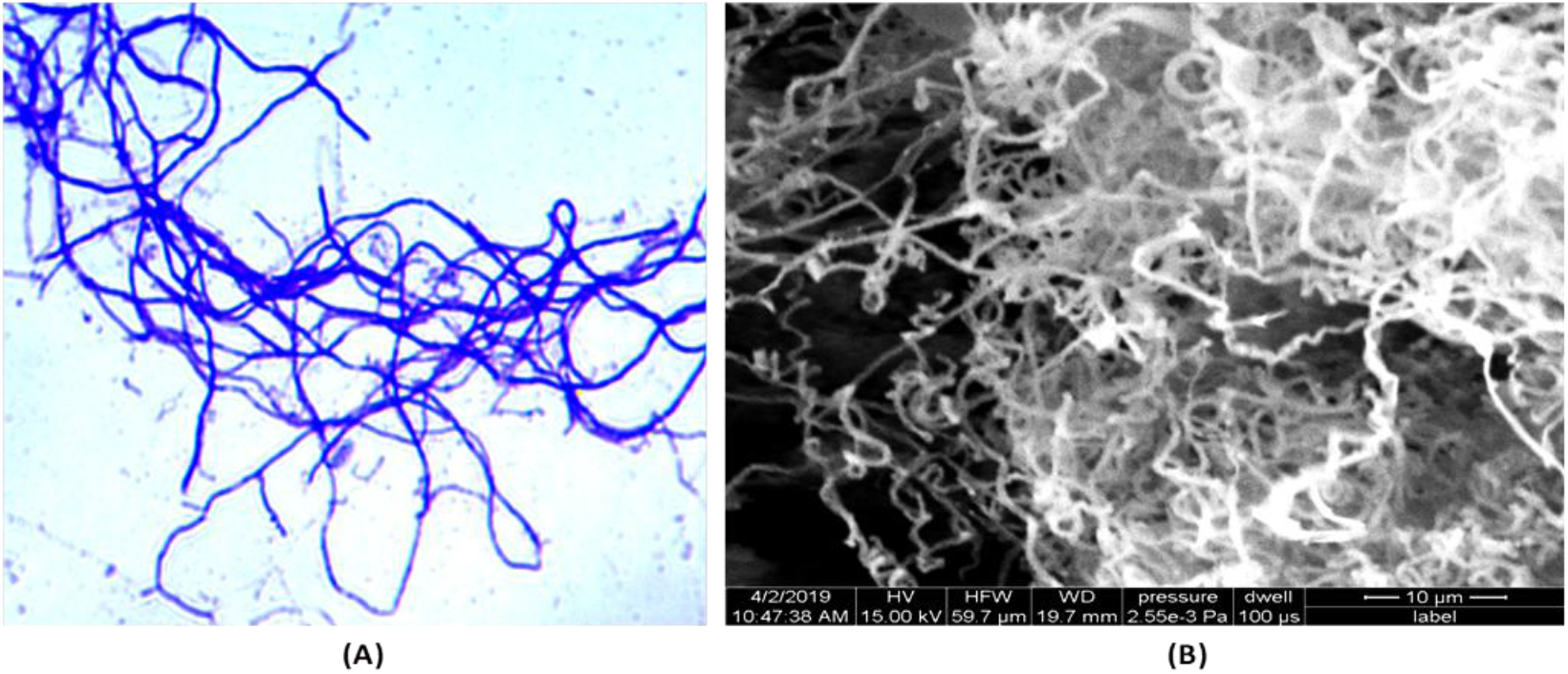
Photomicrograph (1000X) of mycelia (A) and scanning electron micrograph of spores (B) of *Streptomyces* isolate JB140.

**Fig. 3.**
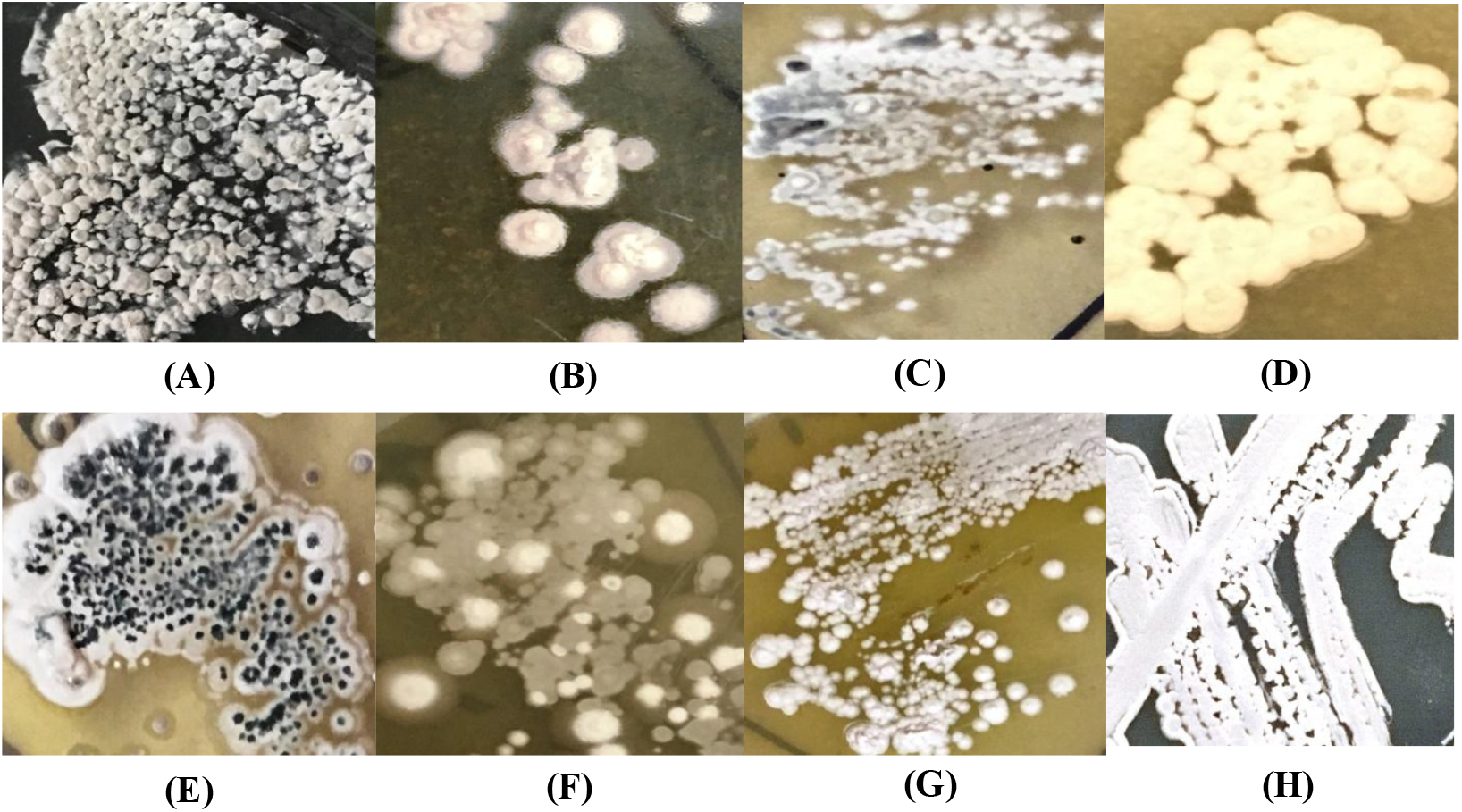
Colonial appearance of the isolate JB140 on (A) modified actinomycetes specific medium (MAS medium) (B) tryptone yeast extract agar (ISP1) (C) yeast extract maltose agar (ISP2), (D) oatmeal agar (ISP3), (E) inorganic salt starch agar (ISP4), (F) glycerol asparagine agar (ISP5), (G) peptone yeast extract iron agar (ISP6), (H) tyrosine agar (ISP7)

### 16SrRNA gene sequence analysis and phylogeny

A comparison of the 16SrRNA gene sequence of the isolate JB140 was made against the available sequences in the GenBank database. It revealed homology of greater than 98% to the members of genera *Streptomyces.* It displayed 97.32% to 98.21% 16SrRNA gene sequence similarities to the closest type strain *Streptomyces* with 98.21% identity to the nearest neighbor *Streptomyces* sp. BN71 (KF479179). The evolutionary relationship of the strain JB140 (MK855143) and the other twenty-six *Streptomyces* spp. has been provided in a phylogenetic tree in figure 4.

**Fig. 4.**
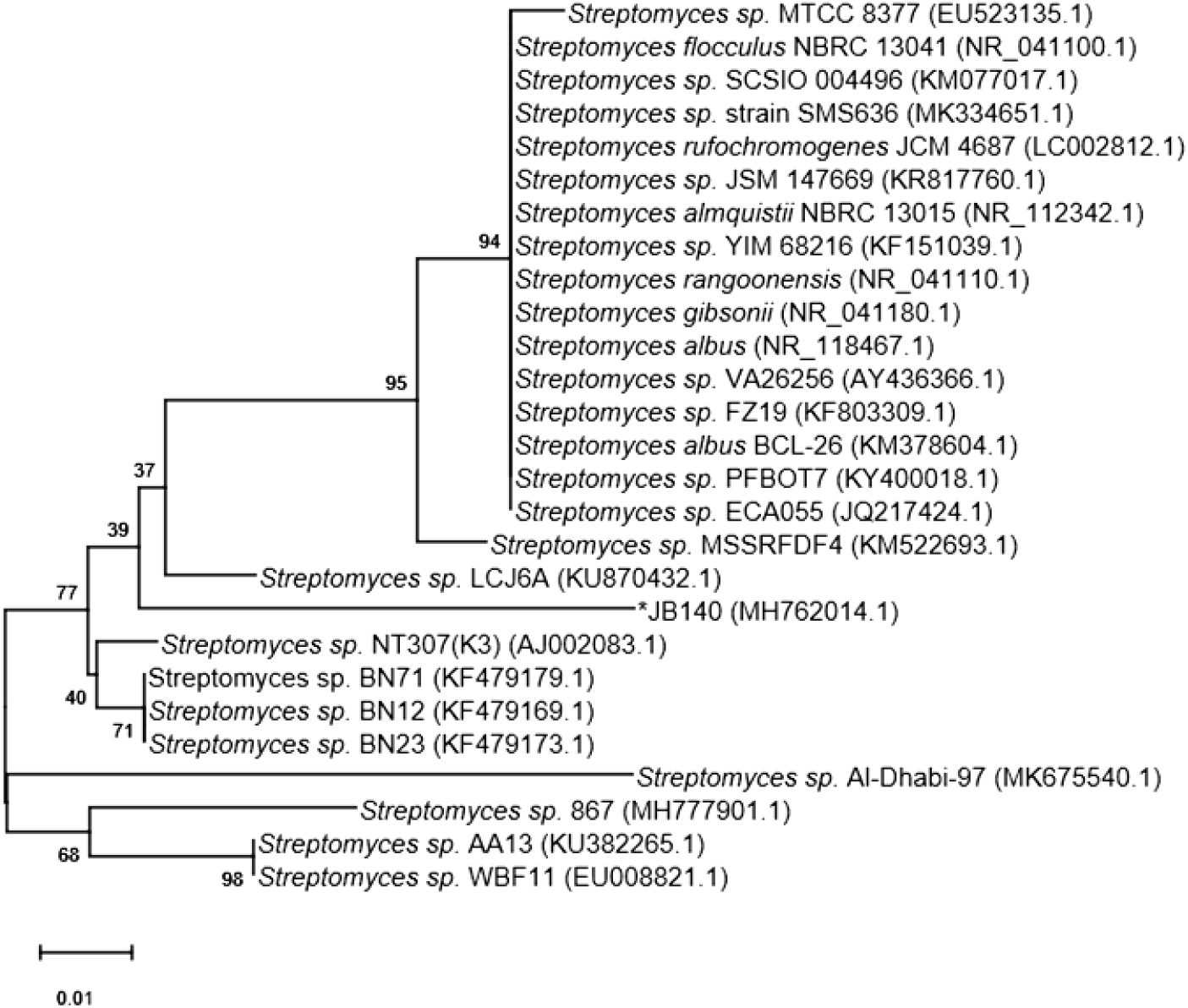
Evolutionary relationship of JB140 was inferred using Neighbor-Joining method. The tree is drawn to scale, with branch lengths in the same units as those of the evolutionary distances used to infer the phylogenetic tree. The evolutionary distances were computed using the Kimura 2-parameter method. The rate variation among sites was modeled with a gamma distribution (shape parameter = 2). Evolutionary analysis was conducted in MEGA X.

### Assessment of antimicrobial activity

The isolate JB140 was analyzed for its antibacterial activity in primary and secondary screening against *Proteus vulgaris, Salmonella typhi* and *Staphylococcus aureus.* The anti-microbial activity of the isolate was estimated by measuring the diameter of the clear zone of growth inhibition (in mm scale). The methanolic extract of the isolate JB140 exhibited weak antimicrobial activity against the pathogenic bacteria tested as the inhibition zone of only 2-3 mm diameter was observed (Fig. 5).

**Fig. 5.**
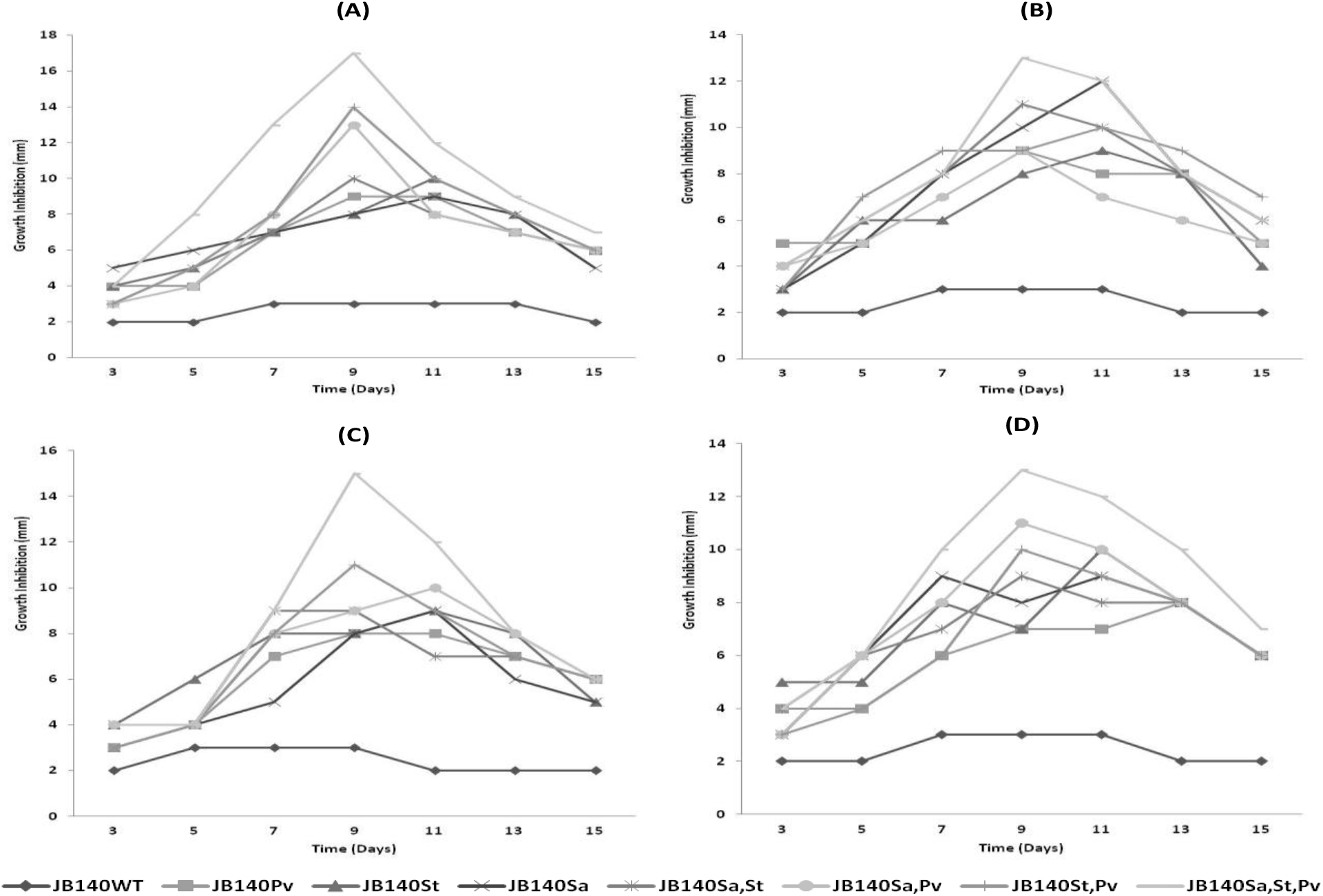
Antimicrobial activity of JB140^WT^ strain and evolved JB140^*Pv*^, JB140^*St*^, JB140^*Sa*^, JBI40^*Sa,St*^, JB140^*Sa,Pv*^, JB140^*St,Pv*^ and JB140^*Sa,St,Pv*^ mutants against *Salmonella typhi* (A), *Staphylococcus aureus* (B). *Proteus vulgaris* (C) and MDR clinical isolate *Escherichia coli* (D)

### Adaptive evolution of *Streptomyces* JB140

Using adaptive laboratory evolution protocol, *Streptomyces* strain JB140^WT^ was competed against three different pathogens in the bi, tri and quadriculture experiments (Fig. 1). In ~45-day long competition experiments, after the third (III) cycle of adaptation-selection (after three serial passages of 10 days each) seven mutants were purified. After each cycle of adaption-selection, the antimicrobial activity of the wild-type and evolved mutants was assayed using the agar well diffusion method. An increased amount of antibiotic production in the evolved mutants was noticed only after the third (III) cycle of adaptation-selection (Fig. 5). Whereas, after the first (I) and second (II) cycles of serial adaptation-selection, a very little or no antimicrobial activity was detected in the evolved mutants. These unique mutants, purified after the third (III) cycle of serial adaptation and selection were designated as JB140^*Pv*^, JB140^*St*^, JB140^*Sa*^, JB140^*Sa,St*^, JB140^*Sa,Pv*^, JB140^*St,Pv*^ and JB140^*Sa,St,Pv*^. All seven mutants produced stable growth inhibitory activity against the tested pathogens. To ascertain that the stimulation of the production of antimicrobials in JB140 mutants is due to three serial adaptive-selection cycles wherein they were exposed to the pathogens, JB140WT (a monoculture experiment without a competing pathogen) was also inoculated using the same procedure. After 45 days (after III cycle in step 7), JB140^WT^ was also assessed for its antimicrobial activity against the tested pathogens. It was observed to be weak and was similar to the antimicrobial activity exhibited by its wild-type counterpart (Fig. 5). Contrary to this, the growth inhibitory effect of the evolved mutants against bacterial pathogens was observed to be from 9mm to 14mm in diameter. The highest antimicrobial activity of the evolved mutants was registered after 9-11 days duration after that interval, it declined. Mutants JB140^*St,Pv*^ and JB140^*Sa,St*^ demonstrated the highest activity against *S. typhi* (Fig. 5A) while mutants JB140^*Sa*^, JB140^*Sa,St*^ and JB140^*Sa,St,Pv*^ were noticed to be highly active against *S. aureus* (Fig. 5B). The highest antagonist effect was observed by JB140^*St,Pv*^ and SB14^*Sa,St,Pv*^ mutants against *P. vulgaris*, (Fig. 5C). Intriguingly, all the evolved mutants were also found to exhibit antimicrobial activity against MDR *E. coli* and the highest activity was registered for the mutants JB140^*St,Pv*^ and JB140^*Sa,St,Pv*^ (Fig. 5D). Furthermore, among seven different competition experiments, the highest antimicrobial activity was observed with the mutants that were evolved against the three pathogens (quadriculture competition experiment) (Fig. 5). The overall observations from the present study suggest that the growth inhibitory effect of the evolved mutants against the tested pathogens including MDR *E. coli* appeared after three serial passages of adaptation-selection only and thus induced antimicrobial activity was not associated with un-evolved, non-competing JB140^WT^ strain. Intriguingly, the whole process of laboratory-based adaptive evolution was reproducible and we could be able to replicate the results when the phenomenon was tested three months apart.

### Characterization of evolved mutants

To confirm that the evolved mutants JB140^*Pv*^, JB140^*St*^, JB140^*Sa*^, JB140^*Sa,St*^, JB140^*Sa,Pv*^, JB140^*St,Pv*^ and JB140*^Sa,St,Pv^* are indeed the true clone of *Streptomyces* JB140^WT^ and are not contaminants, the identity of these were confirmed using 16SrRNA gene sequence analyses. All seven mutants JB140^*Pv*^ (MK850143), JB140^*St*^ (MK852169), JB140^*Sa*^ (MK852177), JB140^*Sa,St*^ (MK852179), JB140^*Sa,Pv*^ (MK852175), JB140^*St,Pv*^ (MK852170), JB140^*Sa,St,Pv*^ (MK852176) were confirmed for their identity as *Streptomyces* similar to their parental wild-type JB140^WT^ (MK855143). Sequence comparison showed that all mutants were true clones of JB140^WT^ strain. The phenotypic characteristics of the evolved mutants were also observed to be the same as their parental JB140^WT^ strain (Fig. 6).

**Fig. 6.**
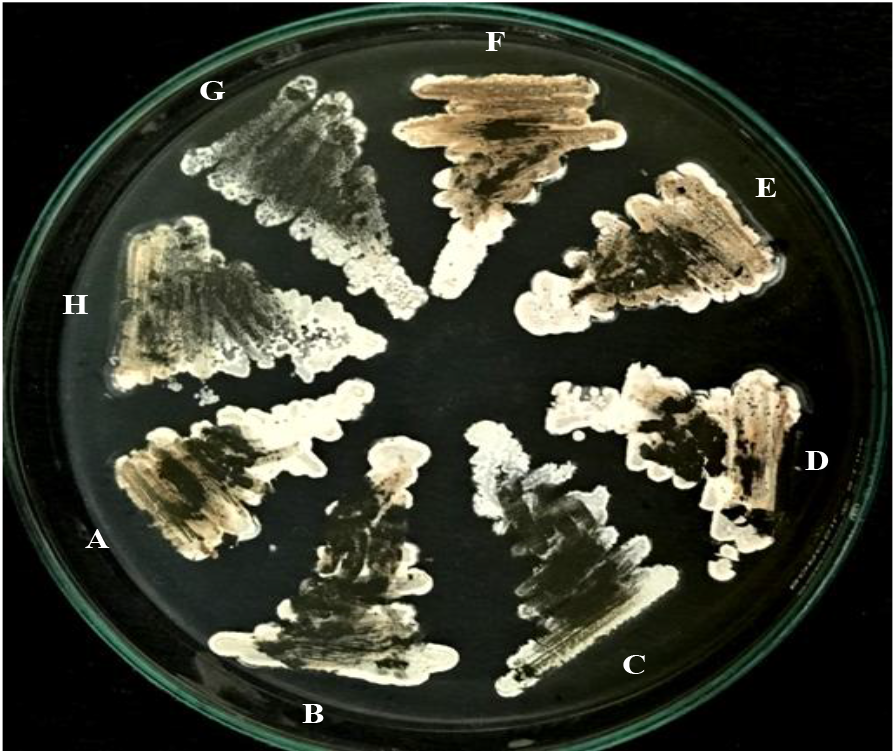
Phenotypic appearance of *Streptomyces* strain JB140^WT^ (A) and its evolved mutants JB140^*St*^ (B), JB140^*Sa*^ (C), JB140^*Pv*^ (D), JB140^*Sa,St*^ (E), JB140^*Sa,Pv*^ (F), JB140^*St,Pv*^ (G) and JB140^*Sa,St,Pv*^ (H) grown on ISP2 medium

### RAPD profiling

In support of adaptive evolution competition experiments and to get evidence of the evolution in actinobacterial JB140^WT^, random amplified polymorphic DNA analysis (RAPD) was performed. Genomic DNA of JB140^WT^ and JB140^*Pv*^, JB140^*St*^, JB140^*Sa*^, JB140^*Sa,St*^, JB140^*Sa,Pv*^, JB140^*St,Pv*^ and JB140^*Sa St,Pv*^ was amplified by using three RAPD primers (1,2 and 3) and compared. The genomic DNA was amplified in separate amplification cycles using RAPD-1, RAPD-2 and RAPD-3 primers individually or in combination (all three). A large number of DNA bands were visualized in the RAPD profiles obtained from the wild-type strain and its mutants that represented significant genetic variations between them (Fig. 7). In a few instances, the RAPD-3 primer was not able to exhibit significant polymorphism between the evolved mutants (Fig. 7C). Nevertheless, detectable differences were observed in the DNA profiles from the wild-type strain JB140 and its evolved mutants using RAPD-1 (Fig. 7A), RAPD-2 (Fig. 7B) and RAPD-1,2,3 primers (Fig. 7D).

**Fig. 7.**
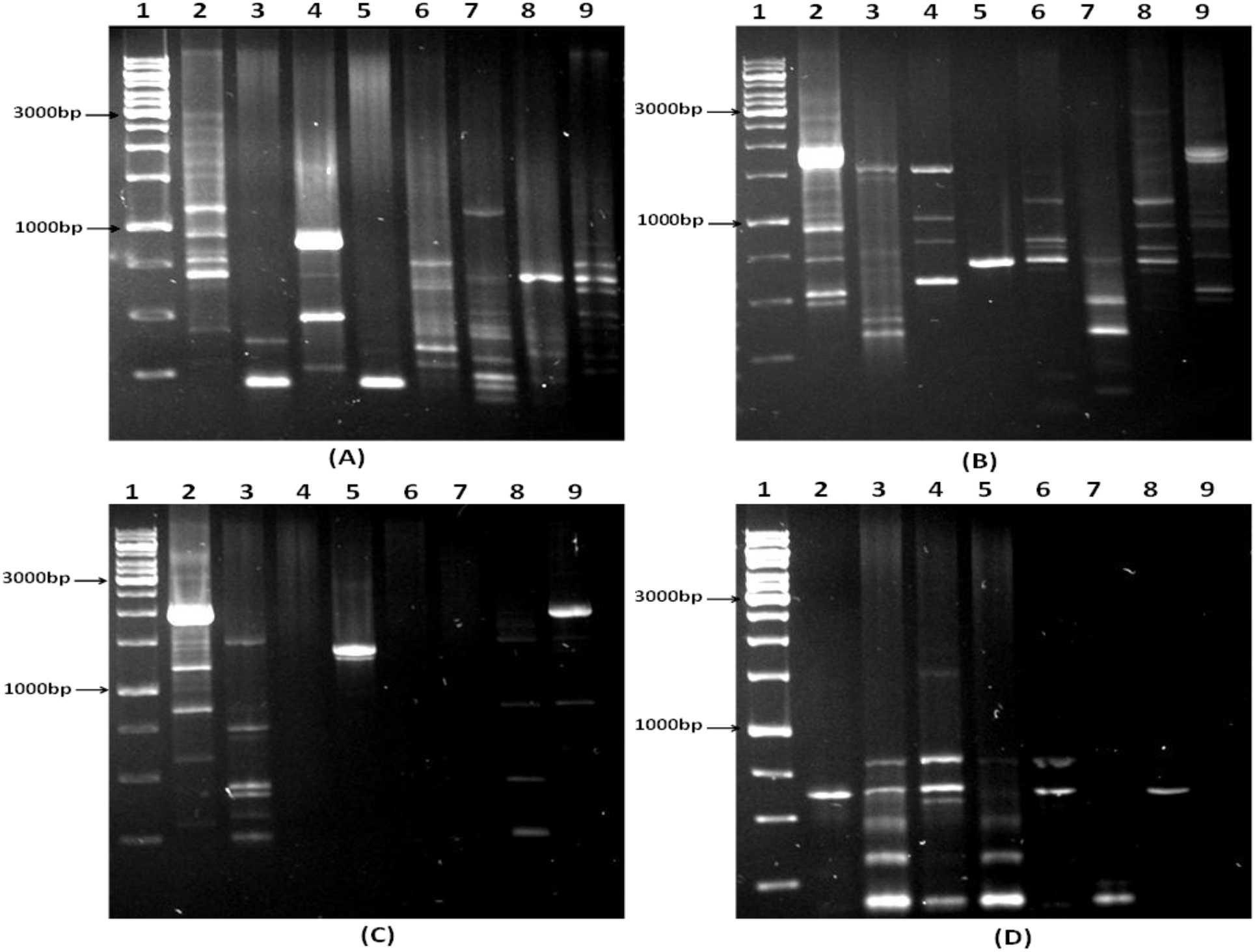
Detection of genetic variation between JB140^WT^ and evolved mutants using RAPD primer 1 (A), RAPD primer 2 (B), RAPD primer 3 (C) and RAPD primers 1,2,3 (D). Lanes 1-1 kb DNA ladder, 2-JB140^WT^, 3-JB140^*Pv*^, 4-JB140^*St*^, 5-JB140^*Sa*^, 6-JB140^*Sa,St*^, 7-JB140^*Sa,Pv*^, 8-JB140^*St,Pv*^, 9-JB140^*Sa,St,Pv*^

## Discussion

Adaptive evolution is considered to be a process in which an organism in culture is grown for generations under some selection pressure to evolve against it. Over a century ago, the experiments involving adaptive laboratory evolution were originally performed (Dallinger 1887; Haas 2000; Bennett and Hughes 2009). However, in the present time, there has been an increase in the number of such experiments that involve microorganisms particularly yeast and bacteria to identify and select evolved phenotypes (Fong and Joyce 2005; Charusanti et al. 2012; Shui et al. 2015; LaCroix et al. 2017). A well-described example of this phenomenon is the development of antibiotic resistance in microbes. Several other examples are also available in literature that well describe this phenomenon such as the evolution of *E. coli* growing in glucose minimal medium to adapt in the media containing glycerol and lactate (Woods et al. 2006; Herring et al. 2006; Barrick et al. 2009; Conrad et al. 2009) and alteration in the productivity of industrially important chemicals (Trinh and Srienc 2009; Hu and Wood 2010). The phenomenon has been tested also in different assays to observe ethanol tolerance (Horinouchi et al. 2010; Stanley et al. 2010; Wang et al. 2011), gene regulation in response to osmotic stress in *Escherichia coli* (Stoebel et al. 2009) and enhanced extracellular electron transfer in *Geobacter sulfurreducens* (Tremblay et al. 2011). We applied this phenomenon in the present work to study the importance of serial passages over time in the adaptive evolution of a weak antibiotic-producing *Streptomyces* strain into a strain exhibiting broad-spectrum antimicrobial activity. In the present study, the pathogenic challenge was presented to the wild-type *Streptomyces* in the biculture, triculture and quadriculture competitions experiments. One of the main findings of the present work is that we were able to detect the evolved phenotypes that could now exhibit strong antimicrobial activity against various pathogens than their parent wild-type strain. Importantly, the evolved mutants also developed the ability to antagonize MDR *E. coli* responsible for UTI infection. Moreover, among seven different competition experiments, the highest antibiotic activity was registered with the mutants that were adaptively evolved against the three pathogens (quadriculture competition). Thus, it could be stated that the evolved unique phenotypes arose only after three serial adaptation-selection cycles and not after initial, short-term exposure to the selection pressure. Other studies have been also performed in which augmented antibiotic production was observed wherein microorganisms were grown in a co-culture (Perry et al. 1984; Sonnenbichler et al. 1984; Slattery et al. 2001; Oh et al. 2007; Kurosawa et al. 2008; Harwani et al. 2018). However, in the present study, the wild-type strain was not detected to exhibit antimicrobial potential until it was serially exposed to bacterial pathogens and passed for three successive adaptation-selection cycles of 15 days each. And importantly, the requirement of three serial passages (three cycles of adaptation-selection) was essential for the wild-type strain to exhibit augmented antimicrobial production against bacterial pathogens.

The pathogens tested in the present study never faced serial passages. It may be speculated that during this period, competitive exclusion (Gause 1932; Hardin 1960) provoked the wild-type *Streptomyces* strain to synthesize chemical molecules to resist the challenge posed by the competing pathogen. When the challenge was presented in the form of more than one pathogen, the higher antagonistic response was elicited by the producer strain. Although the type of chemical molecules (antimicrobials) stimulated and the precise regulatory mechanism of transforming a weak antibiotic-producing strain into a strong one using this phenomenon is yet to be confirmed. But it is important to note, that a simple process as described herein the present study certainly could assist in the discovery of yet undiscovered novel natural metabolites. In the present study, three RAPD primers were also tested individually and in combinations to detect the genetic variations. RAPD profiles yielded significant polymorphism between the wild-type strain and its evolved mutants, but the exact mutational events that occurred during this evolution are yet not known. In this regard, future work will be important to reveal the underlying mechanism responsible for the phenomenon known as adaptive laboratory evolution. To apply and test this method on other systems, a detailed prototype has been also provided in the supplementary data of the manuscript.

## Conclusion

A weak antibiotic-producing *Streptomyces* strain (wild-type) when competed against target pathogens using adaptive evolution protocol led to the identification of seven unique mutant phenotypes. These improved phenotypes (mutants) acquired the ability to constitutively exhibit increased antimicrobial activity against various bacterial pathogens as compared to their parental wild-type counterpart. Moreover, the molecular evidence of the genetic variation collected using RAPD profiling revealed significant polymorphism in the evolved mutants. The constitutive overproduction of the antimicrobials in the mutants was possibly due to the expression of genes that were silent or not expressed in the wild-type parent. Conclusively. the present study signifies that the adaptive laboratory evolution is an efficient tool to select improved phenotypes that can overproduce target-dependent bio-active metabolites. The present study also provides a framework to design more improved selection methods to find out the possible solutions to the issues posed by increasing antimicrobial resistance and provide much-needed novel and alternative antimicrobial therapy.

## Supporting information

Supplementary Material DH et al. Adaptive Evolution

