## Supplementary Material DH et al. Adaptive Evolution for "Adaptive laboratory evolution triggers pathogen-dependent broad-spectrum antimicrobial potency in *Streptomyces*"

### Supplementary data

The following are the supplementary data to this article:

#### Method details

*Background*

The present method describes adaptive laboratory evolution based completion experiments to stimulate antibiotic production in weak antibiotic-producing *Streptomyces* strain ($Producer^{WT}$). To select improved (adaptively evolved) phenotypes, the seven different competition experiments, utilize three serial passages (three cycles of adaptation-selection of 15 days each) of $Producer^{WT}$ against one (biculture; $Producer^{WT}$+Pathogen1 or Pathogen2 or Pathogen3) or two (triculture; $Producer^{WT}$+Pathogen1+Pathogen2 or Pathogen1+Pathogen3 or Pathogen2+Pathogen3) or three (quadriculture; $Producer^{WT}$+Pathogen1+Pathogen2+Pathogen3) target pathogen/s. The present method also describes RAPD profiling of the $Producer^{WT}$ and its evolved mutant phenotypes to observe the molecular evidence of laboratory evolution.

*Material*

- An antibiotic-producing *Streptomyces* strain (here it is referred as a wild-type)
- Type strains of bacterial pathogens or clinical isolates (here these are refereed as three different competitors or targets. MDR bacterial pathogens can also be used)
- ISP-2 (International *Streptomyces* Project) growth medium: yeast extract 4.0 g, malt extract 10.0 g, dextrose 4.0 g, agar 20.0 g, distilled water 1000.0 ml, pH 7.2
- Ethyl acetate
- Methanol
- Bacteriological incubator shaker
- Spectrophotometer ($OD_{600}$) for monitoring cell growth
- Thermocycler for DNA amplification
- Cooling centrifuge
- Taq DNA polymerase
- Genomic DNA (5ng/µl)
- dNTPs (2mM each of dATP, dCTP, dGTP and dTTP)
- MgCl2 (25mM)
- Buffer for DNA polymerase (10mM Tris Hcl, 50mM KCl, pH 8.3)
- Tris-EDTA buffer
- SET buffer (75mMNaCl, 25mM EDTA, 20mMTris, pH 7.5)
- 10-mer OPERON random oligonucleotide primers (5µM): RAPD-1 5'AAGAGCCCGT3', RAPD-2 5'GTTTCCGCCC 3' and RAPD-3 5'GAGGCCCTTC 3'

- Ethidium bromide solution (10mg/ml)
- DNA length marker
- Loading buffer
- Electrophoresis grade agarose
- 1X TBE buffer
- Horizontal gel electrophoresis apparatus
- Gel documentation system
  Note: All chemicals used in the present method are either laboratory or analytical grade. The growth medium, plasticware and glassware were sterilized before use.

*Procedure*
*Adaptive laboratory evolution*

1. Take a loopful of Producer$^{WT}$ (*Streptomyces*) mycelia from glycerol stock to inoculate 8 (eight) different 250mL flasks containing 50mL sterilized ISP2 broth medium. After inoculation, cover the flasks with cotton plugs and incubate at 37$^{o}$C in a shaker incubator at 100rpm for 48h. The first flask inoculated with Producer$^{WT}$ (monoculture) will serve as an internal control (Fig. S1).
2. After observing the visible growth of Producer$^{WT}$, in flasks 2, 3 and 4, inoculate exactly 100 microlitres ($OD_{600}$ =0.50) of different target pathogens 1,2 and 3 respectively (bi-culture competitions) as shown in figure S1. The flasks 2, 3 and 4 will represent the bi-culture competition experiments consist of Producer$^{WT}$ and Pathogen1 or Pathogen2 or Pathogen3.
3. At the same time, in flasks 5,6 and 7 (48h grown culture of Producer$^{WT}$), inoculate exactly 100 microlitres ($OD_{600}$=0.50) of the target pathogens 1+2, 1+3 and 2+3 respectively (tri-culture competitions). The flasks 5, 6 and 7 will represent tri-culture competition experiments consist of Producer$^{WT}$ and Pathogen1+Pathogen2 or Pathogen1+Pathogen3 or Pathogen2+Pathogen3.
4. Similarly, in flask 8, inoculate exactly 100 microlitres ($OD_{600}$=0.50) of the target pathogens 1+2+3 (quadri-culture competition). The flask 8 will represent quadri-culture competition experiment consist of Producer$^{WT}$ and Pathogen1+Pathogen2+Pathogen3.
5. Incubate all eight flasks at 37$^{o}$C for ten days.
6. In the next step, on day 11, spread 100 microlitres of ten days old mono, bi, tri and quadri cultures on the fresh ISP2 agar medium (Fig. S1).
   *Note*: The eight different Petri plates containing the ISP2 agar medium should be spread evenly under aseptic conditions using a laminar bench.
7. Incubate all the inoculated Petri plates at 37$^{o}$C for 5 days to purify Producer$^{WT-}$I (control), Mutant$^{P1}$-I (bi-culture), Mutant$^{P2}$-I (bi-culture), Mutant$^{P3}$-I (bi-culture), Mutant$^{P1+P2}$-I (tri-culture), Mutant$^{P1+P3}$-I (tri-culture), Mutant$^{P2+P3}$-I (tri-culture), Mutant$^{P1+P2+P3}$-I (quadri-culture) as shown in the step 2 in figure S1.
   *Note*: Since the phenotypic appearance of Producer$^{WT-}$I is distinct relative to the bacterial pathogens, hence the colony-forming unit of Producer$^{WT-}$I can be easily picked and purified by streak plate method.
8. After the first cycle of adaption-selection, assay the antimicrobial activities of the wild-type and evolved mutant phenotypes (Producer$^{WT-}$I (control), Mutant$^{P1}$-I , Mutant$^{P2}$-I, Mutant$^{P3}$-I, Mutant$^{P1+P2}$-I, Mutant$^{P1+P3}$-I, Mutant$^{P2+P3}$-I and Mutant$^{P1+P2+P3}$-I) against the

target bacterial pathogens (here pathogen1, pathogen2 and pathogen3) using agar well diffusion method.

*Note:* To evaluate the antimicrobial activity, extract the fermentation broths of the wild-type and evolved mutant phenotypes from their respective flasks with ethyl acetate. The extract should be concentrated to dryness and then dissolve the residue crude extract in methanol. The extracts can be used to assess antimicrobial activity or can be stored at $4^oC$ until further use.

9. The purified mutant phenotypes can be stored here in 20% glycerol (v/v) stocks at -$40^oC$.
10. In the II cycle of adaptation-selection, inoculate eight flasks containing freshly prepared sterilized ISP2 medium again with the purified wild-type and evolved mutants from step 8. After the appearance of visible growth, inoculate these flasks with the target pathogens as described in steps 2-4. Here the different experiments in the II cycle of adaptation-selection will be denoted as experiment 1: Producer$^{WT-}$I (control), experiment 2: Mutant$^{P1}$-I+Pathogen1 (bi-culture), experiment 3: Mutant$^{P2}$-I+Pathogen2 (bi-culture), experiment 4: Mutant $^{P3}$-I+Pathogen3 (bi-culture), experiment 5: Mutant$^{P1+P2}$-I+Pathogen1+Pathogen2 (tri-culture), experiment 6: Mutant$^{P1+P3}$-I+Pathogen1+Pathogen3 (tri-culture), experiment 7: Mutant$^{P2+P3}$-I+Pathogen2+Pathogen3 (tri-culture), experiment 8: Mutant$^{P1+P2+P3}$-I+Pathogen1+Pathogen2+Pathogen3 (quadri-culture) (Fig. S1).

    *Note*: In this step, the flasks should be introduced with freshly grown pathogen/s from their respective stocks.
11. Incubate all eight flasks at $37^oC$ for another ten days (step 3; II cycle).
12. After incubation, purify the mixed cultures (spread and streak plate method as denoted in step **7**) to obtain Producer$^{WT-}$II (control), Mutant $^{P1}$-II , Mutant $^{P2}$-II, Mutant $^{P3}$-II, Mutant $^{P1+P2}$-II, Mutant $^{P1+P3}$-II, Mutant $^{P2+P3}$-II and Mutant $^{P1+P2+P3}$-II (step 4; Fig. S1).

    Note: After the II cycle of adaption-selection, as denoted in step 8, assay the antimicrobial activities of the wild-type and evolved mutant phenotypes against the target pathogens.
13. The purified mutant phenotypes-II can be stored here in 20% glycerol (v/v) stocks at -$40^oC$.
14. In the III cycle of adaptation-selection, inoculate eight flasks containing freshly prepared sterilized ISP2 medium again with the purified wild-type and evolved mutants from step **12**. After the appearance of visible growth, inoculate these flasks with the target pathogens as described in steps 2-4. The different experiments in the III cycle of adaptation-selection will be denoted as experiment 1: Producer$^{WT-}$II (control), experiment 2: Mutant $^{P1}$-II+Pathogen1 (bi-culture), experiment 3: Mutant $^{P2}$-II+Pathogen2 (bi-culture), experiment 4: Mutant $^{P3}$-II+Pathogen3 (bi-culture), experiment 5: Mutant $^{P1+P2}$-II+Pathogen1+Pathogen2 (tri-culture), experiment 6: Mutant $^{P1+P3}$-II Pathogen1+Pathogen3 (tri-culture), experiment 7: Mutant $^{P2+P3}$-II Pathogen2+Pathogen3 (tri-culture), experiment 8: Mutant $^{P1+P2+P3}$-II Pathogen1+Pathogen2+Pathogen3 (quadri-culture) to finally receive the purified wild-type Producer$^{WT-}$III and evolved Mutant $^{P1}$-III, Mutant $^{P2}$-III, Mutant $^{P3}$-III, Mutant $^{P1+P2}$-III, Mutant $^{P1+P3}$-III, Mutant $^{P2+P3}$-III, Mutant $^{P1+P2+P3}$-III from their respective mono,bi,tri, quadri-culture competition experiments using spread-streak plate method as described in steps 6,7 and 12 and illustrated in figure S1 (step 5 and 6).

15. After the III cycle of adaption-selection, assay the antimicrobial activities of the wild-type and evolved mutant phenotypes (Producer$^{WT-}$III (control), Mutant $^{P1}$-III , Mutant $^{P2}$-III, Mutant $^{P3}$-III, Mutant $^{P1+P2}$-III, Mutant $^{P1+P3}$-III, Mutant $^{P2+P3}$-III and Mutant $^{P1+P2+P3}$-III) against the target bacterial pathogens (here pathogen1, pathogen2 and pathogen3) or MDR pathogens using agar well diffusion method (step 7).
    *Note:* To evaluate the antimicrobial activity, extract the fermentation broths of the wild-type and evolved mutant phenotypes with ethyl acetate. The extract should be concentrated to dryness. Dissolve the residue crude extract in methanol and store at 4$^{o}$C until further use.
16. The purified mutant phenotypes can be stored in 20% glycerol (v/v) stocks at -40$^{o}$C.

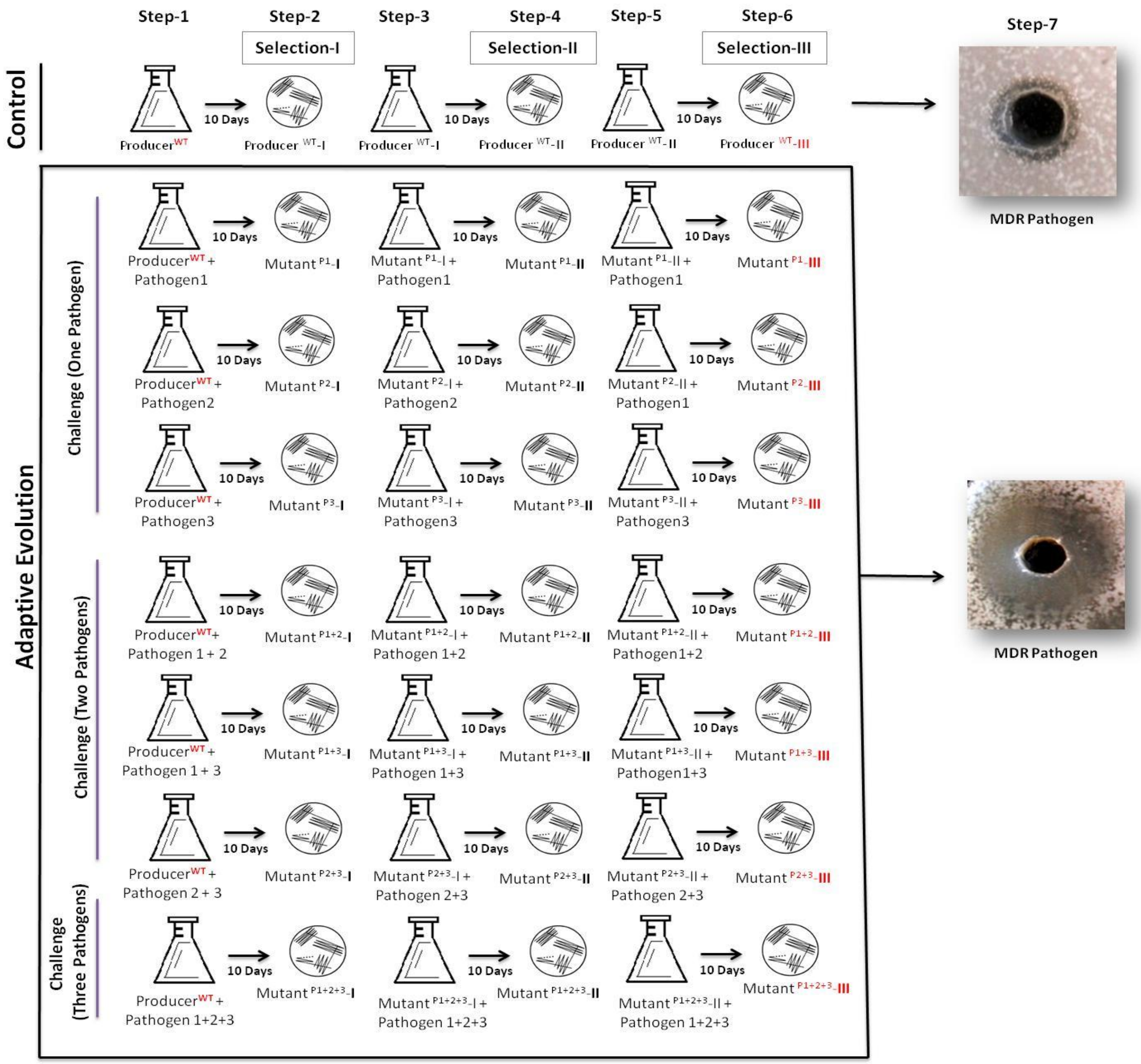


**Fig. S1** Adaptive evolution protocol (seven-steps) wherein a wild-type producer strain is competes against bacterial pathogens to stimulate antibiotic production

*Random amplified polymorphic DNA (RAPD) profiling*

1. Use 10ml of purified cultures (grown at 40±1$^{o}$C; $OD_{600}$=0.7) of wild-type producer and mutant phenotypes in centrifuge tubes. Centrifuge these at 8000 rpm for 5 minutes. Discard the supernatant carefully and collect the cell pellets in 2.0 ml Eppendorf tubes individually.
2. Wash pellet with TE buffer (Tris-EDTA) and re-suspended in 0.6 ml SET buffer (75mMNaCl, 25mM EDTA, 20mMTris, pH 7.5).
3. Add lysozyme to a concentration of 1mg ml$^{-1}$ and incubate at 45$^{o}$C for 1hour. Add 0.1 volume 10% SDS and 0.5mg ml$^{-1}$ proteinase K and incubate at 55$^{o}$C with occasional inversion for 2.5 hours.
4. Add one-third volume 5M NaCl and 1 volume chloroform and incubate it at room temperature for 0.5 hours with frequent inversion. Centrifuged the mixture at 10000rpm for 15 minutes and transfer the aqueous phase to a fresh tube using a pipette tip (blunt-ended).
5. Precipitate the chromosomal DNA by adding one volume 2-propanol (invert it gently), wash it twice with 70% ethanol, dry in speed-vacuum and dissolve in 100µl of pure water.
6. Treat the dissolved DNA with 10mg ml-1 RNaseA at 37$^{o}$C for 30 minutes.

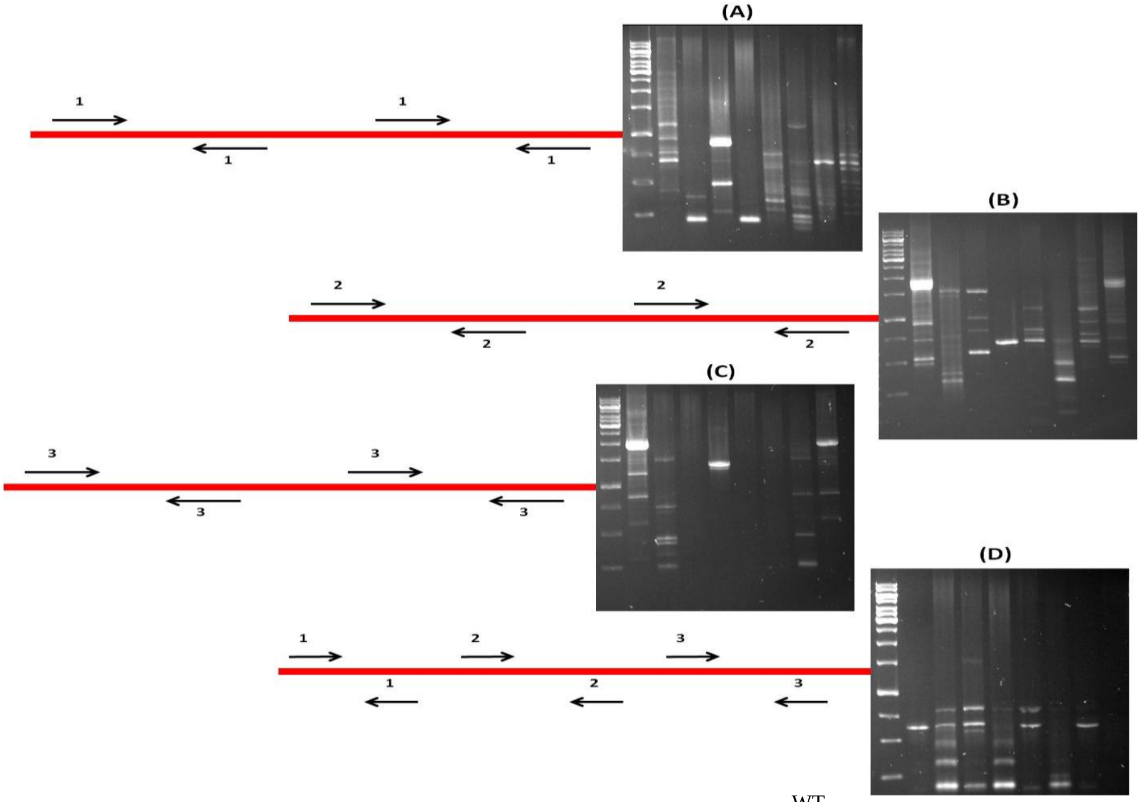


**Fig. S2** RAPD analysis to detect genetic variation between Producer$^{WT}$ and evolved mutants using RAPD primer 1 (A), RAPD primer 2 (B), RAPD primer 3 (C) and RAPD primers 1,2,3 (D). Lanes in the agarose gel denote 1 kb DNA ladder followed by polymorphism between the wild-type producer strain and its mutant phenotypes selected after the III cycle of adaptation-selection in the method

7. Extract the DNA in the same volume of phenol/chloroform/isoamyl alcohol (25:24:1) and precipitate with 2.5 volume ethanol (cold) and 0.1 volume 3M sodium acetate.
8. Wash the pellets using 70% ethanol, dry and dissolve in 100µl TE or pure water.
9. Visualize DNA in electrophoresis using 2% agarose gel and reveal it subsequently using ethidium bromide staining.
10. For RAPD reactions, prepare a master mix (for seven mutant phenotypes and control (wild-type producer)) that contains all the above components. Use three RAPD primers (RAPD-1 5'AAGAGCCCGT3', RAPD-2 5'GTTTCCGCCC 3' and RAPD-3 5'GAGGCCCTTC 3') individually (RAPD-1 or RAPD-2 or RAPD-3) or in combination (all three; RAPD-1+ RAPD-2 +RAPD-3) in the four different separate amplification cycles (Fig. S2) using a DNA thermocycler. Each 20µl reaction volume will contain the following:

| S.No. | Master mix ingredients | Volume (microlitres) | Final Concentration |
|---|---|---|---|
| 1 | Genomic DNA | 5.0 | 25ng |
| 2 | Buffer (10mM Tris Hcl, 50mM KCl, pH 8.3) | 2.0 | 1X |
| 3 | dNTPs | 1.0 | 0.2mM |
| 4 | Each Primer | 1.0 | 0.5µM |
| 5 | Taq DNA polymerse | 0.2 | 1 unit |
| 6 | MgCl2 | 1.0 | 3mM |
| 7 | Purified water to make | 20 | |

11. Reactions without DNA will serve as a negative control.
12. Programme the thermocycler for an initial denaturation step of 5 minutes at 95°C, followed by 40 cycles of DNA denaturation at 95$^{o}$C for 1 minute, primer-annealing at 32$^{o}$C for 1 minute and extension cycle at 72$^{o}$C for 1 minute, with a final extension at 72$^{o}$C for 10 minutes.
13. After amplification, prepare four 2% (w/v) agarose gels in 1X TBE buffer containing ethidium bromide (0.5µg/ml). Add 3µl of the loading dye to each sample.
14. Load the DNA length marker and the different eight RAPD reactions (PCR amplified) into the wells of their corresponding gels. Run the gels in 1X TBE buffer at 50V.
15. Visualize the gels under UV transilluminator or gel documentation system and record photographs (Fig. S2).
